# Habitat Suitability Modeling and Ecological Forecasting of Northern Goshawk Nesting Habitat

**DOI:** 10.1101/2022.02.04.479095

**Authors:** Marylin. E. Wright, Jenelle. Jackson, Risto. Tornberg, Erica. Higa, A. Clayton, Sean. McCartney, Dustin H Ranglack, Nate Bickford

## Abstract

Northern Goshawks (*Accipiter gentilis*) are dynamic forest raptors often used as a management indicator species in forest management planning. Despite their use as indicators, there is a limited understanding of habitat requirements for goshawks, especially in dry forested landscapes as climate change occurs. We examined goshawk nesting habitat on the Helena-Lewis and Clark National Forest (HLCNF) in central Montana to further understand climate change on goshawk nesting habitat. The HLCNF receives low annual precipitation, making this forest useful to understand climate change in dry forest in structure and ecology. We measured characteristics of nest sites throughout the HLCNF and used principle component analysis to predict important variables for nest trees and nesting habitat for goshawks in this landscape. We also used remote sensing and GIS to create a composite map predicting areas of high, medium and low suitability for nesting habitat for goshawks in the HLCNF. We used the composite map to predict potential impacts of mountain pine beetle (*Dendroctonus ponderosae*), fire, and climate change to Northern Goshawk habitat in the HLCNF. The best indicators of a chosen nest tree were diameter at breast height, tree height, and tree status, which can be considered indicative of trees preferred for nesting by goshawks. The best indicators for nest forest were canopy closure, slope angle, and slope position, which indicate habitat areas of high suitability for goshawk nesting. Two forecast models where developed based on climate change predictions. After overlapping a projected mountain pine beetle risk map and fire frequency risk map onto the modeled of high suitability goshawk nesting habitat, we calculated that 52 % of modeled high suitability habitat may be at risk for mountain pine beetle blight and 66 % high suitability habitat at risk for frequent wildfires, which will reduce goshawk habitat quality.

## Introduction

Northern Goshawks (*Accipiter gentilis*) are the largest *Accipiter* and are found throughout the Holarctic region. Because of this species’ important role as an apex predator in boreal and temperate forest ecosystems, Northern Goshawks have become a major focus of many forest management plans (Malengo et al. 2013, Stein Foster et al. 2010, Lehr 2014). Established under the U.S. National Forest Management Act in 1982, Management Indicator Species have been a required component of national forest management plans. Species selected as management indicators were intended to help focus forest management decisions, provide a method for assessing forest biodiversity and health, and serve as the principal monitoring species during forest plan implementation (USFS 1982).

Despite their use as indicator species, there is a limited understanding of the habitat requirements for Northern Goshawks (Squires and Reynolds 1997). Previous studies have demonstrated that goshawks hunt in predominantly forested areas and nest in mature forests where there is high canopy cover and an open understory (Reynolds et al. 1982, Speiser and Bosakowski 1987, Hayward and Escaño 1989, Siders and Kennedy 1994, Squires and Ruggiero 1996, Squires and Reynolds 1997, Youtz et al. 2008, Greenwald et al. 2005). However, in much of the United States nest site locations are limited due to their preference for moderately sloped areas that are north to east facing (Squires and Reynolds 1997). Many of the studies that have examined nest stand and nest tree requirements for goshawks have focused on areas of wet forest that receive substantial annual precipitation (Hayward and Escaño 1989, Lilieholm et al. 1993, Bull and Hohmann 1994, Squires and Ruggiero 1996, Bruggeman et al., 2011). The areas of the HLCNF used in this study identifying goshawk nesting habitat represent predominantly warm, dry forest with little annual precipitation (max ~330 mm;(DeBlander 2002, Wright et al., 2019). Although other studies of goshawk habitat on dry forest have been performed, this study will endeavor to identify possible changes to the forest via climate change (Beck et al., 2011,,.;’ Youtz et al., 2008, Keane, Morrison, and Fry, 2006, Greenwald et al. 2005). Due to the low precipitation, forest ecology on the HLCNF is much different than that of a wet forest located at higher elevations and climate change will have different impacts to the forest. Precipitation has a direct effect on forest structure at all levels, and forest structure has been implicated as one of the primary determinants of high suitability goshawk nesting habitat (Erickson 1987, Rissler 1995, Reynolds 2004, Greenwald et al. 2005, Youtz et al. 2008).

Changes in climate have exacerbated habitat loss caused by both pine beetle (*Dendroctonus ponderosae*) kill and forest fires (Kurz et al. 2008). Pine Beetle is an indigenous North American insect with major impacts on forests in western North America due to loss of environmental controls of the population (Kurz et al. 2008). Warmer temperatures and variation in seasonality during recent decades have allowed the mountain pine beetle (*Dendroctonus ponderosae*) to proliferate in national forest lands throughout Western North America, creating large areas of disturbance dead pines that may not support nesting goshawks (Kurz et al. 2008). Warming trends and earlier springs combined with large deposits of downed woody vegetation have also allowed wildfires to increase in frequency, size, and intensity (Westerling et al. 2006). Surveys conducted by the U.S. Forest Service indicated that goshawks in the Jefferson Division of HLCNF have been impacted due to climate-related expansion of mountain pine beetle (Murphy 2010). Climate change also presents another set of challenges to goshawk survival, in that extreme weather conditions and seasonal variation can affect goshawk egg and fledgling survival because cold, wet springs and delayed snowmelt at higher elevations can delay egg laying (Squires and Kennedy 2006).

By focusing on the HLCNF, the goal of this study was to characterize goshawk nesting habitat in a xeric western forest, and predict how future disturbance (pine beetles and increase fire frequency as well as intensity) will influence goshawk nesting habitat. Our first objective was to characterize nest tree and nest habitat at known nest sites throughout the study area to determine the variables most indicative of goshawk nesting habitat. These variables, along with ancillary data, informed our model by use of NASA Earth observations as inputs for habitat suitability modeling. The next objective was to create and analyze three different habitat suitability models and form a composite map to identify potential nesting habitat for goshawks in the HLCNF. The final objective was to forecast the impacts of future climate change on nesting habitat, potential mountain pine beetle encroachment, and fire risk by the year 2050.

## Study Area

The HLCNF encompasses ~7,300 km^2^ in central Montana. The forest is separated into two management areas: the Jefferson and Rocky Mountain Divisions. The ~4,900 km^2^ Jefferson Division is further divided into five mountain ranges: Castles, Crazies, Highwoods, Little Belts, and Snowies. Many of these geographic areas are discrete geologic units with unique landforms, vegetation, cultural histories, and recreation opportunities. Elevation ranges from 1,370 to 2,850 m. Primary forest types consist of Douglas fir (*Pseudotsuga menziesii*), lodgepole pine (*Pinus contorta*), ponderosa pine (*Pinus ponderosa*), and aspen (*Populus tremuloides*); all of which are common nest trees for goshawks Reynolds 2004, Greenwald et al. 2005, Youtz et al. 2008. The U.S. Forest Service oversees forest management within the study area, including timber, fire management, and recreational activities (USDA Forest Service 2018, HLCNF).

## Methods

### Field Collection

We used the protocol outlined by the Northern Goshawk Inventory and Monitoring Technical Guide to derive a sampling strategy (Woodbridge and Hargis 2006). To determine presence of goshawks and locate a nest stand, we used a combination of search surveys and broadcast acoustical surveys along transects. Our survey method consisted of visual confirmation of goshawk presence including nests, whitewash on trees, prey remains, and molted feathers around the nest tree (Woodbridge and Hargis 2006; Victor Murphy personal communication). We also broadcasted previously taped goshawk calls on a digital amplifier at variable decibels along transects to elicit a defensive response from nesting goshawk pairs (Kennedy and Stahlecker 1993, Woodbridge and Hargis 2006). Once the presence of a goshawk pair was determined, we located the nest site by following the goshawk back to its nest and recorded that location with a handheld GPS. Nest sites were visited at least twice in a given year between June and August. Nest site locations were all combined into a historic database from surveys in the 1980’s. Surveys were ongoing for 8 years (2007 – 2013) for 190 nest site locations. These surveys were conducted as part of a U.S. Forest Service survey project.

We recorded values for a list of variables for each nest site pertaining to the physical habitat characteristics of both the tree containing the nest and the surrounding area. We recorded the species of nest tree, status of the nest tree (dead or alive), diameter at breast height (DBH), nest height, and nest tree height. For the area surrounding the nest site, we recorded placement of the nest (elevation, nest orientation, slope aspect, slope angle, slope position (summit, shoulder, footslope, toeslope)). As well as habitat characteristics (age class of the trees, overstory canopy closure, evidence of previous timber cutting (yes or no), distance to a water source, tree species present within the immediate area around the nest (10 m radius around the nest), and vegetation cover within the immediate area around the nest).

#### Data Analysis

This study examined different variables that affect goshawk nest selection. We examined variation in nest tree and nest habitat variables using principal component analysis without rotation (Janžekovič and Novak 2012). For the nest tree, we used five variables in our analysis including tree species, status (dead or alive), DBH, nest height, and nest-tree height. Tree status was coded as a dichotomous variable with zero indicating the nest tree was dead and one indicating the nest tree was alive. Tree species were coded by rank with the most common nest tree species having the higher number (Douglas fir coded as 4, lodgepole pine coded as 3, quaking aspen coded as 2, ponderosa pine coded as 1). All other variables were entered as continuous variables—DBH, nest height, nest-tree height.

We used six variables to assess characteristics of habitat surrounding nest sites including elevation, canopy closure, overstory canopy species, slope aspect, slope angle, age class of trees, and slope position. Age class of the trees was coded by rank with the most common age class present at nest stands, mature (coded 2) and a mixture of mature and pole trees (coded 1). Age class was determined with standard USFS protocol (Victor Murphy personal communication).

### Habitat Suitability Modeling

#### Data Acquisition

We used Northern Goshawk nesting sites recorded from 1985 to 2013 as presence data in our models of habitat suitability. The historic sites were visited in 2007 and any nest sites that were located on radically changed landscape (clear cut, etc.) were not used in the analysis. We assessed these datasets for redundancy by removing overlapping nest sites within a 5 m buffer to account for GPS error, and combined them into one set of 190 nesting sites.

The U.S. Forest Service maintains a large geospatial library containing data for national forests in the western U.S. (ID, WA, MT, ND, and SD). We utilized this library to obtain data on vegetation characteristics within the HLCNF and to derive a set of nesting habitat variables for Northern Goshawks. The U.S. Forest Service Vegetation Map (VMap) and Forest Inventory and Analysis (FIA) programs survey, analyze, and produce geospatial data layers for various forest attributes. We used the VMap product to obtain vegetation metrics in vector format for tree canopy cover-class size. Tree canopy cover is classified by percent canopy cover. We derived basal area, in m^2^/ha, from the FIA product at 250 m resolution. We also derived vegetation height from the Existing Vegetation Height (EVH) layer of the U. S. Geological Survey (USGS) Landscape Fire and Resource Management Planning Tools (LANDFIRE) program. This layer provides an average height of the dominant vegetation at 30 m resolution.

Using USGS Earth Explorer, we acquired Shuttle Radar Topography Mission (SRTM) 1 arc-second (30 m) void-filled data, which were used to derive terrain elevation, slope, and aspect. We acquired Landsat 8 Operational Land Imager (OLI) data from the USGS Earth Resources Observation and Science (EROS) Center as three separate tiles for the dates of 7 September 2014 and 2 August 2015. We used these tiles to create a forest cover map for inclusion in habitat suitability modeling. We acquired precipitation data from Precipitation Measurement Missions, Tropical Rainfall Measuring Mission (TRMM), and Global Precipitation Measurement (GPM), for the months of February to June 2010 to 2015. TRMM data were acquired as a level 3 product (3B43) at 0.25-degree spatial resolution and one-month temporal resolution. GPM data were acquired as a level 3 product (3IMERGM) at 0.1-degree spatial resolution and one-month temporal resolution. We downloaded both data sets from the Science Team On-Line Request Module (STORM), a web-based data access interface hosted by NASA Goddard Space Flight Center.

#### Data Processing

We created data layers for each environmental variable to model habitat suitable for Northern Goshawk nesting habitat. These variables included elevation, slope, aspect, precipitation, land cover, vegetation height, canopy cover, and basal area. We included variables based on their having been identified in previous research as being related to goshawk nest sites. Five individual SRTM elevation tiles spanning the study area were acquired and topographic layers for slope and aspect derived from this 30 m DEM. VMap data were converted to a 30 m raster format to create separate layers for canopy cover class. FIA data for basal area were resampled from 250 m to 30 m spatial resolution to match the spatial resolution of the other environmental variables. TRMM and GPM data were resampled from 0.25-degree spatial resolution, and 0.1-degree spatial resolution respectively, to 30 m spatial resolution. Precipitation was then averaged over the nesting season and used as an environmental input into each model. For creating a land cover map of the study area, three tiles of Landsat 8 data were sorted together. Each data layer was subset to match the spatial extent of the HLCNF, Jefferson Division.

#### Data Analysis

After pre-processing the Landsat 8 OLI data for Top of Atmosphere (TOA) reflectance values, further analysis was carried out to create a land-cover classification for the HLCNF using bands 2-7; the results of which were two categories: forest and non-forest. Three separate hard classifications were undertaken using different methods to create land cover maps to model habitat suitability. Maximum likelihood and segmentation classifications were carried out using TerrSet software (Mahmoud and Divigalpitiya 2017), and Monte Carlo unmixing and subsequent hard classification was carried out using CLASlite software (Asner 2009). Once all classifications were completed, 200 random samples were generated as an ArcGIS shapefile (Esri ArcGIS v.10.3) and converted to a KML point file. These randomly generated points were then mapped in Google Earth and verified as being either forest or non-forest. The resulting text file was used as ‘ground truth’ and converted to a raster file in a GIS where an accuracy assessment was performed with each hard classification. The object-based classification (segmentation) resulted in the highest overall accuracy rate of 88 % and was subsequently used to develop habitat suitability models.

We utilized three presence-only modeling software (Maximum Entropy, Mahalanobis Typicality, and BioMapper) to construct a habitat suitability model for Northern Goshawk nesting habitat. The empirical models determine the environmental constraints of the species by relating locations where the Northern Goshawk was observed to different environmental factors. We ran the Maximum Entropy (Maxent) and Mahalanobis Typicality models through TerrSet’s Habitat and Biodiversity Modeler application (Mahmoud and Divigalpitiya 2017). Maxent is a widely used modeling software that estimates the probability distribution of a species using a maximum entropy approach, where the expected value of each environmental variable matches the empirical average (Phillips et al. 2010). Mahalanobis Typicality expresses the likelihood that a set of environmental variables at a specific location is typical of the known location of the species, or that the species distribution is normal with respect to environmental gradients (Hernandez et al. 2008). The stand-alone software, BioMapper 4.0, was used for the third model. BioMapper is based on Ecological Niche Factor Analysis (ENFA), which computes the environmental factors that most explain the ecological distribution of a species (Hirzel et al., 2002). The following eight environmental variables were input into each model: elevation (m), slope (percent), aspect (direction of slope), land cover (forest/non-forest), vegetation height (m), tree canopy cover class (percent cover), precipitation (mm), and basal area (m^2^/hectare). A correlation matrix was first computed with all environmental variables to evaluate the correlation of the variables that will be used for running each model (Table 1). Any two variables with a correlation coefficient greater than 0.7 were flagged. The variables tree size and tree canopy were highly correlated at 0.987, therefore tree size was excluded and the remaining eight variables were used in each model.

**Table 1.** Correlation matrix for environmental variables selected for habitat suitability modeling

| **COR. MATRIX** | **Aspect** | **DEM** | **Basal area** | **Land cover** | **Precipitation** | **Slope** | **Tree canopy** | **Tree Size** | **Veg height** |
| --- | --- | --- | --- | --- | --- | --- | --- | --- | --- |
| **Aspect** | 1 | -0.016 | 0.038 | -0.04 | -0.009 | 0.037 | 0.027 | 0.027 | 0.017 |
| **DEM** | -0.016 | 1 | 0.142 | -0.004 | 0.307 | 0.027 | 0.143 | 0.147 | -0.118 |
| **Basal area** | 0.038 | 0.142 | 1 | -0.543 | 0.114 | 0.148 | 0.071 | 0.146 | 0.301 |
| **Land cover** | -0.04 | -0.004 | -0.543 | 1 | -0.054 | -0.021 | -0.034 | -0.121 | -0.336 |
| **Precipitation** | -0.009 | 0.307 | 0.114 | -0.054 | 1 | 0.061 | 0.047 | 0.062 | -0.026 |
| **Slope** | 0.037 | 0.027 | 0.148 | -0.021 | 0.061 | 1 | 0.218 | 0.232 | -0.025 |
| **Tree canopy** | 0.027 | 0.143 | 0.071 | -0.034 | 0.047 | 0.218 | 1 | 0.987 | -0.188 |
| **Tree Size** | 0.027 | 0.147 | 0.146 | -0.121 | 0.062 | 0.232 | 0.987 | 1 | -0.134 |
| **Veg height** | 0.017 | -0.118 | 0.301 | -0.336 | -0.026 | -0.025 | -0.188 | -0.134 | 1 |

Once the 3 models were run using the 8 environmental variables, each model was validated by generating a percentage of random evaluation sample points and re-running each model. For this study 75% of the total presence points were randomly generated. Each model was then trained with this percentage of points and the rest of the observations were used to test each model. Results of modeling with evaluation sample points were then tested for accuracy and Relative Operating Characteristic (ROC) were generated and area under the curve (AUC) values interpreted for ranking the accuracy of each model.

Based on the results of each habitat suitability model for goshawk nesting sites, an average-value, composite map was created showing a gradation of probabilities for nesting sites in the study area. The composite map was generated by calculating the mean pixel value from all three habitat suitability models. Equal interval classification of the average-value map was used to display areas of high, medium, and low suitability nesting habitat.

### Ecological Forecasting

Data were acquired for climate projections from the Goddard Institute for Space Studies ModelE/Russell Model (GISS-E2-R) for three representative concentration pathways (RCPs). Each RCP describes possible global climate scenarios based on greenhouse gas concentration trajectories adopted by the Intergovernmental Panel on Climate Change (IPCC) for its fifth Assessment Report (IPCC 2014).

Mountain pine beetle infestation projections were downloaded from the U.S. Forest Service 2012 National Insect and Disease Risk Maps/Data (“National Insect & Disease Risk Map” 2019). Raster data measuring total projected basal area loss in square meter per hectare through 2027 were used. Mean fire return interval data were downloaded from the USGS LANDFIRE program displaying historical fire regimes by categories of varying temporal length (years).

The mountain pine beetle and mean fire return interval data were resampled to 30 m spatial resolution and georeferenced to NAD UTM zone 12N. The mountain pine beetle risk map was reclassified so each pixel with projected basal area loss (range, 0.09 m^2^/hectare - 19.8m^2^/hectare) was considered at risk. A fire frequency risk map was created by combining the following mean fire return interval classes: 11-15 years, 16-20 years, 21-25 years, 26-30 years, and 31-35 years. This risk map identifies areas that could be affected by natural wildfire by 2050.

As disturbances such as beetle outbreaks and wildfire adversely affects the forest health, and therefore goshawk nesting habitat, we forecasted what areas of the HLCNF may be viable for nesting based on the current conditions and future scenarios. To assess how bioclimatic conditions will differ in the future, we compared current average precipitation and minimum temperature data to the projected data extracted from the GISS-E2-R climate model for RCP 2.6, RCP 4.5, and RCP 8.5. Additionally, we overlaid and compared the mountain pine beetle risk map and the fire risk map with areas of high suitability nesting habitat as determined by modeled results.

## Results

### Nest-Tree Characteristics

We located 190 nest sites. Many of these nests were in the same territories and some of the nest were old and falling apart. We analyzed 23 total goshawk nests, both active regular territory (*n* = 15) and inactive regular territory (*n* = 7 active the year before during surveys; unknown: *n* = 1 irregular territory and has been abandoned); however, four of these sites were present on private property, so we were unable to take all descriptive measurements at these locations (three sites on the Rocky Mountain front, one in the Little Belt Mountains). Whereas the 23 nests only reflect a small subset of known nests on the forest, time constraints prevented the field crew from taking measurements at all nest sites. Over half of the nest sites were found in Douglas fir (*n* = 12) (Fig. 2). Only two nest sites were observed in dead trees, one on the Rocky Mountain front and one in the Little Belt Mountains. The mean diameter of the 19 recorded nest trees was 29.4 cm 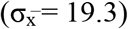, and the mean tree height was 17 m (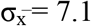; Table 2). Nest trees ranged from 18 to 78 cm in diameter, but the average nest trees measured from 25 to 35 cm. Tree height ranged from 43 to 70 m, but the average trees measured from 55 to 65 m.

**Figure 1.**
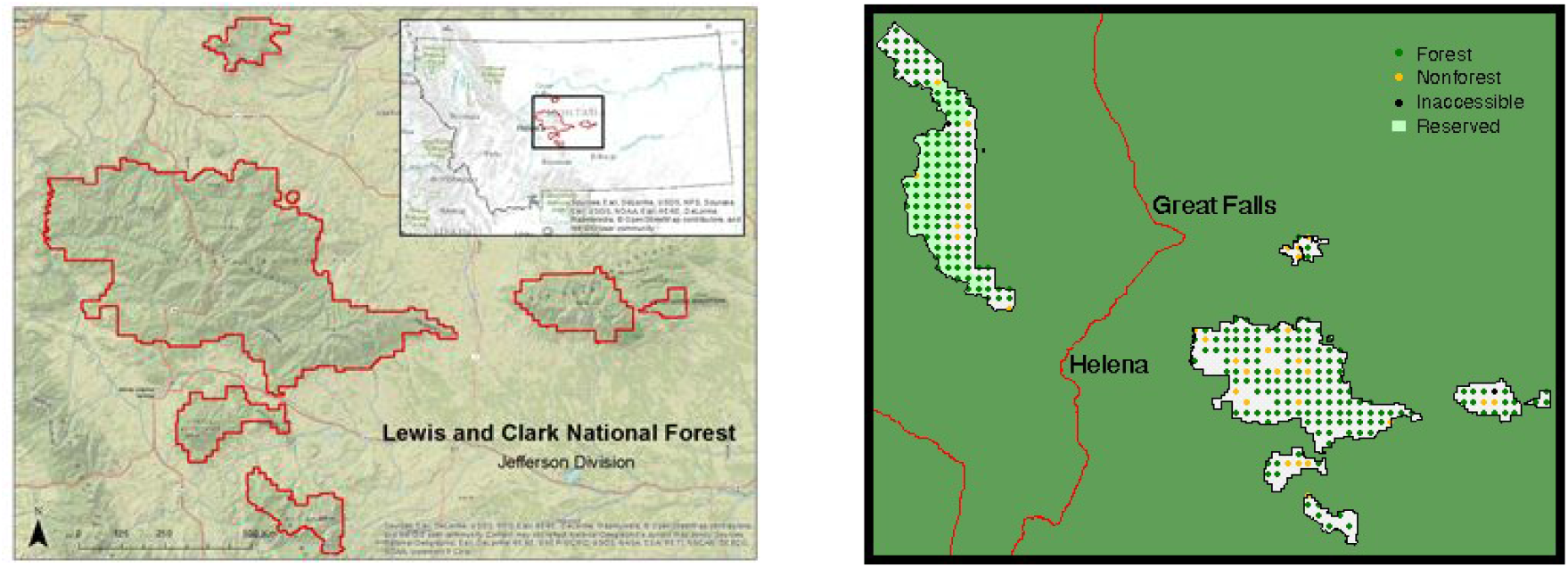
Forest cover in the HLCNF Rocky Mountain and Jefferson Divisions (left) used for vegetation analysis (USDA 1995) and the Jefferson Division (right) used for habitat modeling

**Figure 2.**
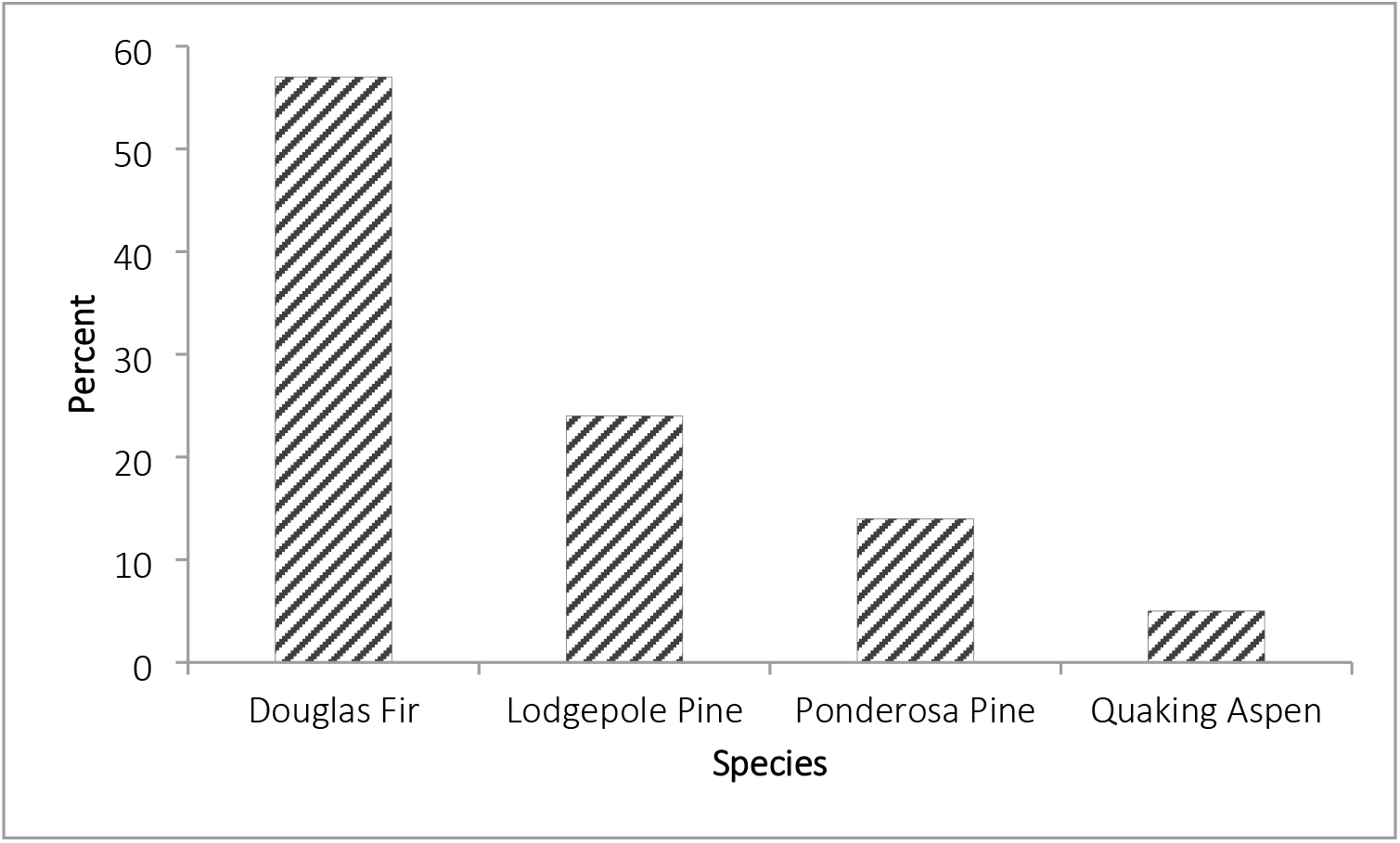
Percentage of each nest tree species

**Table 2.** Mean and standard deviation of diameter at breast height (DBH), nest height, and tree height for all nest trees across the study area

| Location | *n* | Tree DBH (cm) |  | Nest Height (m) |  | Tree Height (m) |  |
| --- | --- | --- | --- | --- | --- | --- | --- |
|  |  | Mean | SD | Mean | SD | Mean | SD |
| Rocky Mountain Front | 10 | 32.6 | 16.0 | 7.6 | 1.4 | 17.0 | 2.6 |
| Little Belts | 6 | 31.8 | 8.2 | 9.5 | 2.5 | 16.3 | 2.2 |
| Castles | 2 | 50.8 | 39.5 | 21.5 | 7.5 | 17.2 | 5.8 |
| Big Snowies | 1 | 27.9 | *NA* | 10.7 | *NA* | 17.4 | *NA* |
| All Ranges | 19 | 34.0 | 16.4 | 9.8 | 4.9 | 16.8 | 2.6 |

Note: DBH, nest height, and tree height measurements were not taken at all nest sites due to private property restriction

Three principal component axes had eigenvalues > 1.0, and together they accounted for 80 % of the total variance. The first principal component (34 % of the total variance) was most highly positively correlated with DBH and tree height. There were no negative correlations. High values on this axis represented trees with greater DBH and total height. The second principal component, which accounted for 26 % of the total variance, was most highly positively correlated with tree status (alive or dead) and most negatively correlated with tree height. The third principal component, which accounted for 20 % of the total variance, was most highly correlated with tree status and most negatively correlated with tree species. The highest positively correlated values on each of the first three principal components (DBH, tree height, and tree status) are considered indicative of trees preferred for nesting by goshawks (Table 3, Fig. 3).

**Figure 3.**
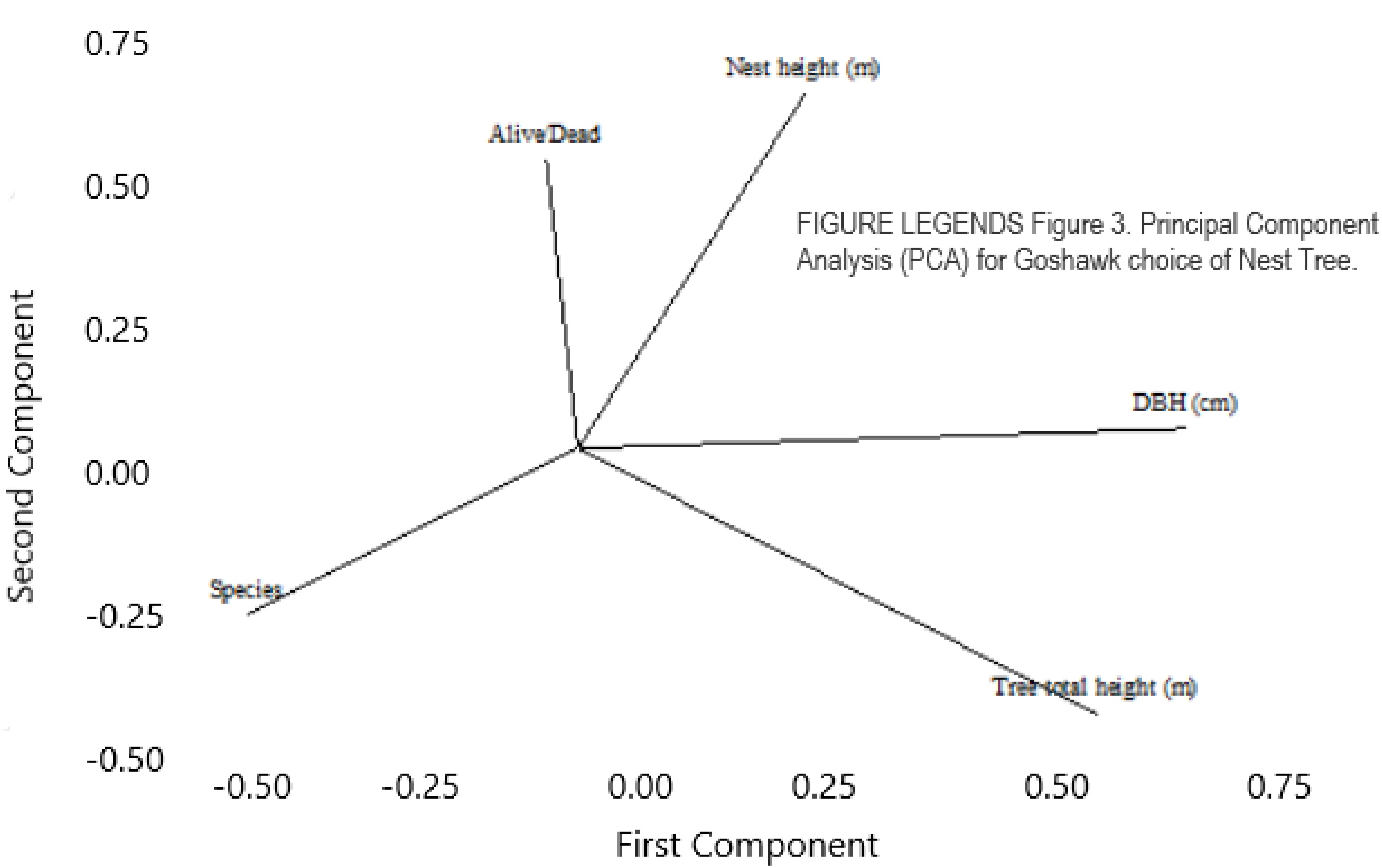
Loading plot for nest tree PC1 and PC2.

**Table 3.** Principal component analysis results of goshawk nesting trees in the study area

| Variable | PC1 | PC2 | PC3 |
| --- | --- | --- | --- |
| DBH | 0.677 | -0.005 | 0.313 |
| Tree Height | 0.526 | -0.525 | 0.170 |
| Species | 0.440 | 0.256 | -0.540 |
| Nest Height | 0.266 | 0.580 | -0.295 |
| Alive/Dead | 0.021 | 0.568 | 0.704 |
| Eigenvalue | 1.68 | 1.32 | 1.01 |
| Proportion | 0.34 | 0.26 | 0.20 |
| Cumulative | 0.34 | 0.60 | 0.80 |

### Nest Habitat Characteristics

For nest stands across all four mountain ranges, the mean elevation was 1728 m with a total range of only 366 m (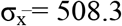). Habitat stands were composed mainly of Douglas fir and lodgepole pine with equal representation of Engelmann spruce and ponderosa pine (17 %) and quaking aspen and subalpine fir (9 %). We observed mature age class trees at all nest sites, though three sites (all on the Rocky Mountain front) had a mix of mature and pole timber. We observed evidence of previous timber cutting at only three nest sites (one on the Rocky Mountain Front and two in the Little Belt Mountains). All other nest habitats showed little-to-no disturbance. Average canopy closure across all mountain ranges was 48 % (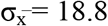). The least amount of canopy closure was observed at the one nest site in the Big Snowy Mountains (15 %) and the highest amount of canopy closure were observed in the Castle Range (75 %). Slope average aspect across ranges was 144° which corresponds to a northwest orientation. Average slope angle was 24° and the most common slope position for nest stands was in the shoulder area. The average distance to the next closest known nest site was 242 m with a range from 25 to 1100 m.

Three principal component axes had eigenvalues > 1.0, and together they accounted for 76 % of the total variance for nest habitat characteristics. The first principal component (35 % of the total variance) was most highly positively correlated with canopy closure and slope angle, and most negatively correlated with slope aspect. High values on this axis represented habitats with increased canopy closure and similar variance in slope angle whereas low values represented little variation in slope aspect. The second principal component, which accounted for 24 % of the total variance, was positively correlated with slope position and most negatively correlated with elevation and age class. This demonstrates a lack of correlation among the nest-stand position on the slope, elevation, and age class of trees in the area surrounding nests. The third principal component, which accounted for 18 % of the total variance, was most highly correlated with slope angle and most negatively correlated slope aspect and slope position. The highest positively correlated values on each of the first three principal components (canopy closure, slope angle, and slope position) are considered indicative of preferred areas for goshawks nesting habitat (Table 4, Fig. 4).

**Figure 4.**
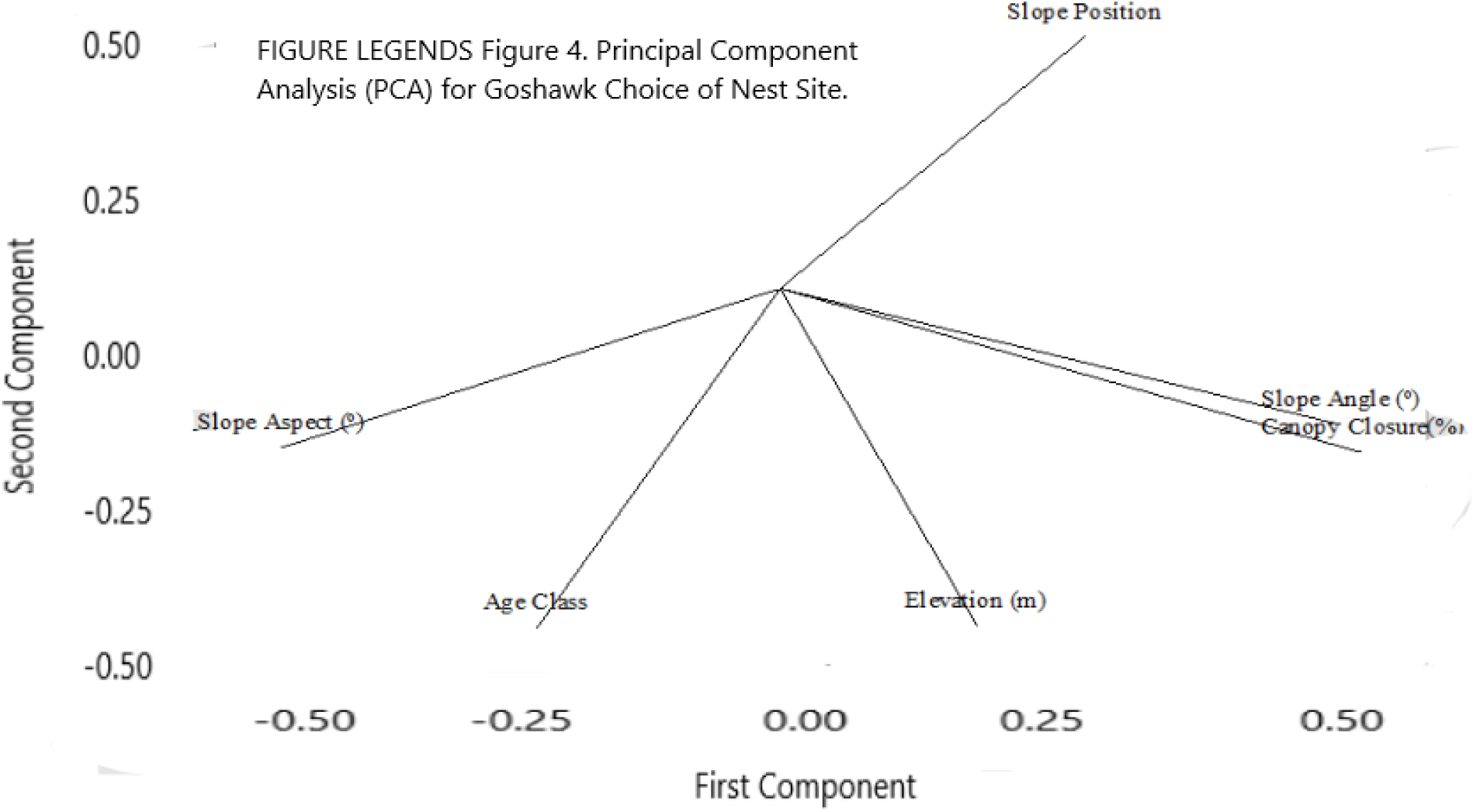
Loading plot for nesting habitats PC1 and PC2.

**Table 4.** Principal component analysis results of goshawk nesting habitats in the study area

| Variable | PC1 | PC2 | PC3 |
| --- | --- | --- | --- |
| Elevation | 0.189 | -0.558 | -0.311 |
| Canopy Closure | 0.557 | -0.272 | -0.32 |
| Slope Aspect | -0.476 | -0.266 | -0.491 |
| Slope Angle | 0.537 | -0.227 | 0.382 |
| Age Class | -0.232 | -0.564 | 0.066 |
| Slope Position | 0.292 | 0.417 | -0.64 |
| Eigenvalue | 2.08 | 1.42 | 1.08 |
| Proportion | 0.35 | 0.24 | 0.18 |
| Cumulative | 0.34 | 0.58 | 0.76 |

### Habitat Suitability Modeling

In the final composite suitability map (Fig. 5), approximately 350 km^2^ (or 7 %) of the Jefferson Division of the Helena-Lewis and Clark National Forest was classified as high suitability nesting habitat for the Northern Goshawk. There are approximately 725 km^2^ of marginal nesting habitat (15 % of the study area), while the remaining 78 % of the HLCNF Jefferson Division was classified as low suitability for goshawk nesting sites.

**Figure 5.**
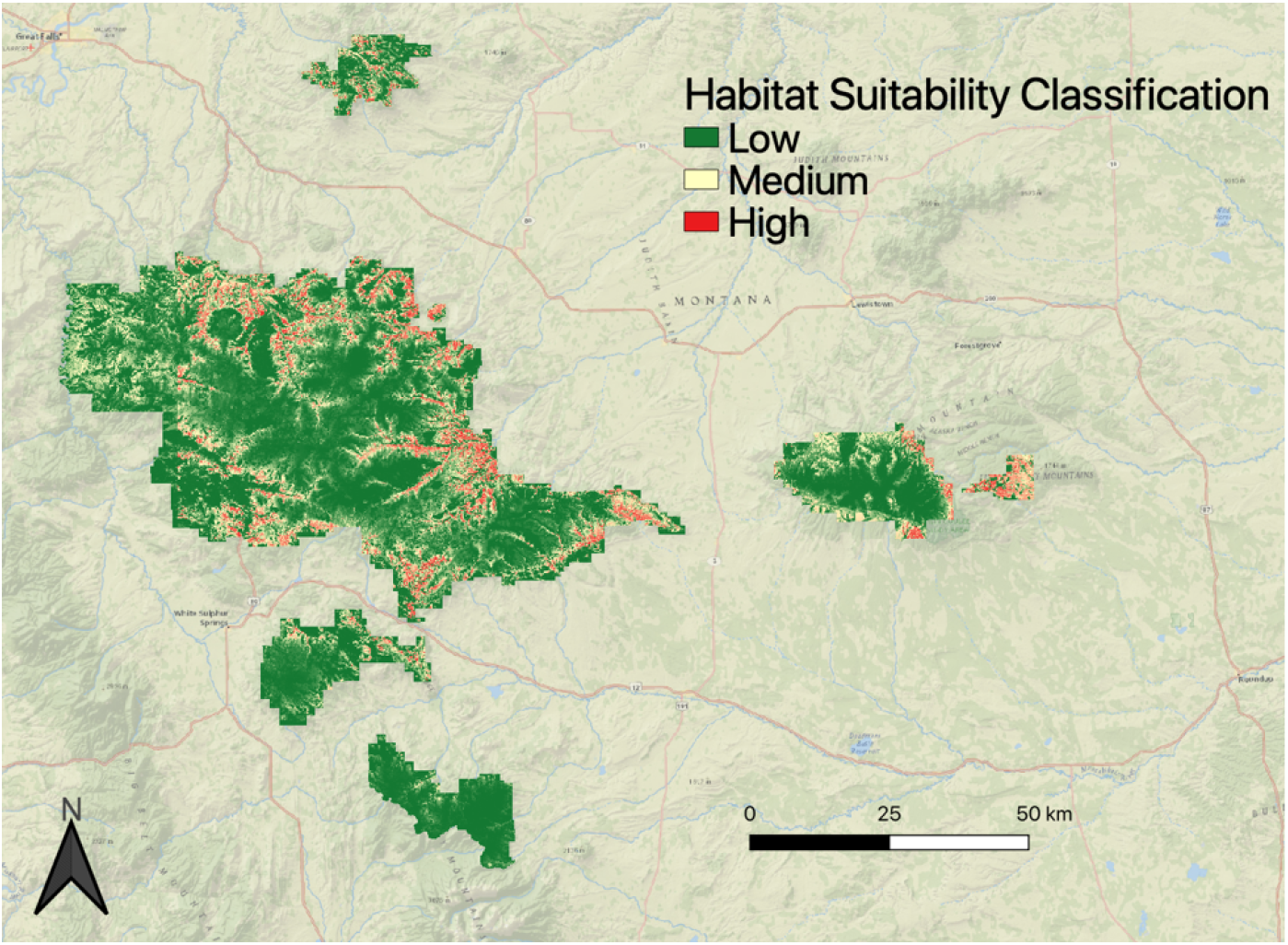
Composite map combining BioMapper, Maxent, and Mahalanobis Typicality habitat suitability maps to show areas of high, medium, and low suitability nesting habitat for northern goshawks within the Helena-Lewis and Clark National Forest, Jefferson Division

**Figure 6.**
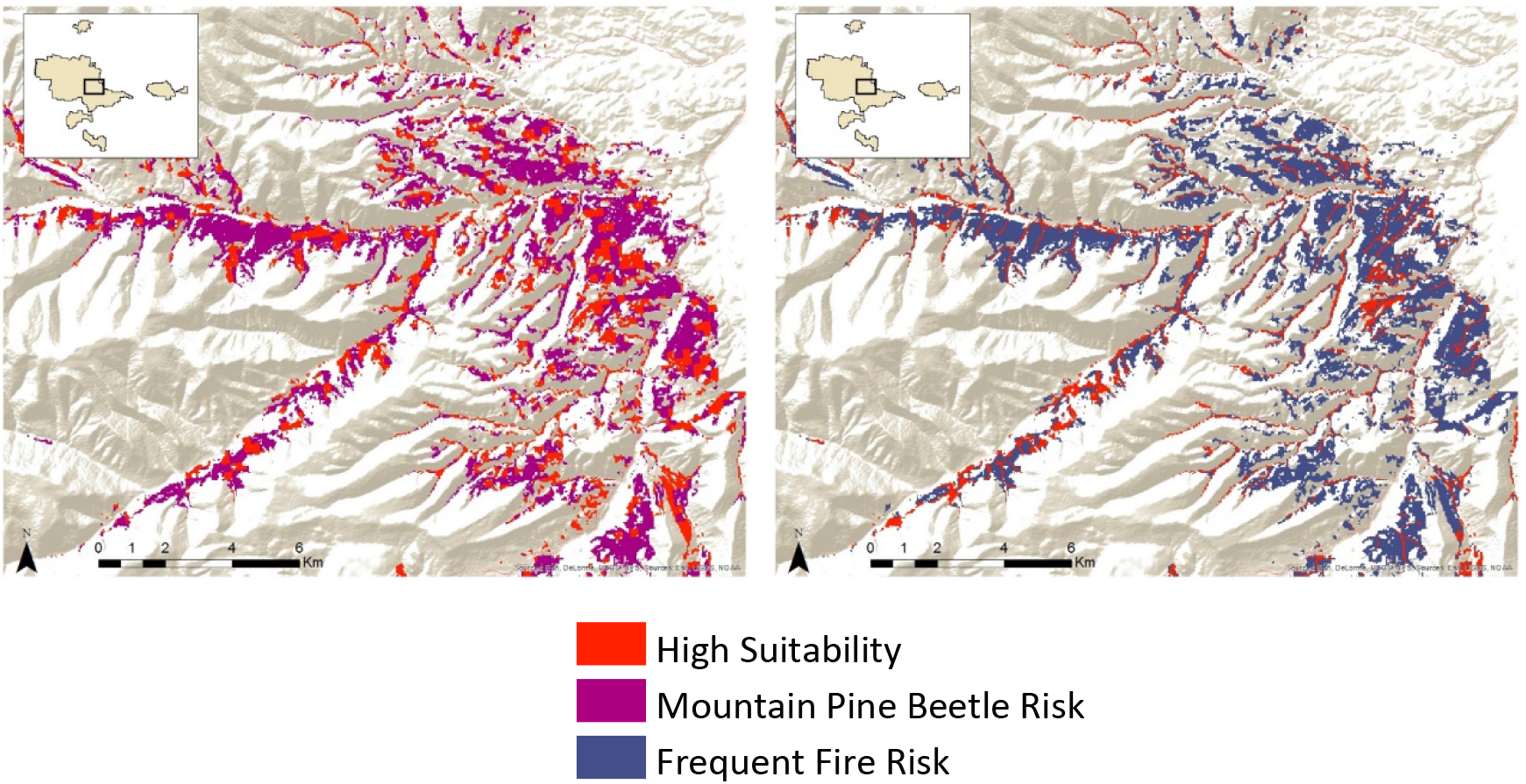
The map on the left shows the amount of high suitability habitat at risk due to mountain pine beetle blight projected to 2027. The map on the right shows amount of high suitability habitat at risk for frequent fires by 2050.

We extracted the habitat suitability values in the composite map based on the 190 nest locations to check the performance of the habitat suitability model (Table 5). One-hundred-three nests coincided with high suitability areas and 66 nests were located in medium suitability areas. There is some uncertainty in the habitat suitability map because not all models are perfect and that 21 nests were found in areas modeled as low suitability. Not all nests were active, and some of the 21 nest areas may have been subject to habitat disturbances between the last known active year and present, which could not be accounted for in our model. Another possible explanation is that 2^nd^ year nesters will sometime choose poor locations for nesting.

**Table 5.** Validation of the composite map by extracting the habitat suitability values on the basis of the 190 nest locations.

|  | *Number of Nest Sites (percentage of total nests, n=190)* |
| --- | --- |
| *High* | 103 (54%) |
| *Medium* | 66 (35%) |
| *Low* | 21 (11%) |
| *Range of suitability values* | 0.022 to 0.986 |
| *Mean suitability* | 0.655 |
| *Median suitability* | 0.698 |

After overlapping the reclassified 2027 projected mountain pine beetle risk map onto the modeled best available goshawk habitat, we calculated that 52% of modeled high suitability habitat may be at risk for mountain pine beetle blight (Fig. 8). We overlaid the reclassified fire frequency risk map and the modeled high suitability nesting habitat map using the “intersect” function and identified 66% of high suitability habitat at risk for frequent wildfires, possibly driven by drying conditions from climate change (Fig. 8).

## Discussion

Through vegetation analysis and habitat modeling, we were able to determine that high suitability goshawk nesting habitat is some of the most underrepresented habitat area available in the HLCNF. We found that DBH, tree height, and tree status (alive or dead) were the best predictors of which nest trees goshawks would use for nesting, whereas canopy closure, slope angle, and slope position were the most important variables for predicting the surrounding habitat characteristics. Similar to studies conducted in other portions of the species range, goshawks in the HLCNF preferred larger trees with a dense distribution and closed canopy (Reynolds et al. 1982; Saunders 1982; Moore and Henny 1983; Hall 1984; Spieser and Bosakowski 1987; Crocker-Bedford and Chaney 1986; Hayward and Escaño 1989; Lang 1994; Siders and Kennedy 1994; Daw 1996; Siders and Kennedy 1996; Squires and Ruggiero 1996; Desimone 1997; Patla 1997; Daw et al. 1998; Keane 1999; Finn et al. 2002). In the HLCNF, only 7% of available habitat matched these specifications.

One of the variations between the vegetation data collected for this study and that of other studies was the mean DBH and height of nest trees and trees within the surrounding habitat. For all mountain ranges within our study area, the mean DBH was 34 cm and the mean nest tree height was 17 m. A study conducted west of our study area on the Rocky Mountain Front found that the average DBH for nest trees ranged from 38.1 to 50.8 cm, with a mean height of 19.7 m (Clough 2000). The low mean DBH and tree height observed in our study is likely directly related to the low annual precipitation on the eastern side of the Rocky Mountain Front and throughout the HLCNF Jefferson Division mountain ranges. With little rainfall, tree growth in the HLCNF is slower overall, leading to a greater density of smaller, shorter trees in mature forest (DeBlander 2002; Anderson et al. 2013).

A greater density of smaller, shorter trees is particularly concerning when considered in conjunction with forecasting outputs showing areas of high suitability goshawk nesting habitat in the HLCNF that will be susceptible to habitat loss in the future. Warmer temperatures are thought to permit mountain pine beetles to extend their range to higher altitudes and latitudes and quicken the beetle’s reproductive and growth cycles (Bentz et al. 2009). Warmer winter and spring seasons and drier conditions will also reduce cold-induced beetle mortality, and thus allow higher populations of pine beetles than in cooler conditions (Bentz et al. 2009). We analyzed the reclassified 2027 projected mountain pine beetle risk map by overlaying the modeled high suitability goshawk nesting habitat with the mountain pine beetle risk map and calculated that 52% of modeled high suitability habitat may be at risk for mountain pine beetle blight (Fig. 8). Continued warming and decreased precipitation is expected to put even greater amounts of habitat at risk in the future as there will likely be an increase in beetle infestation in areas that are not currently affected (Kurz et al. 2008).

Warmer and drier climate will also create conditions that make forested areas more susceptible to wildfires and possibly increase the frequency of fires. We observed the reclassified fire frequency risk map and overlaid the modeled high suitability habitat with this map and found that 66% of modeled high suitability habitat may be at risk for frequent wildfires (Fig. 8). It is, however, not guaranteed that wildfires will occur at the modeled high suitability goshawk nest sites because wildfires effect the whole landscape not just high suitability goshawk nest sites. Regardless, warming trends will possibly contribute to continued goshawk habitat loss in the HLCNF.

The limiting factor that could not be accounted for in this project was exact location of the nesting habitat that goshawks prefer to validate the suitability model. Although we utilized eight environmental variables that influence goshawk nesting habitat in our habitat modeling, there is some uncertainty in our results. We were not able to incorporate an important requirement for nesting habitat in our modeling; goshawks prefer to nest in forested areas that have a closed canopy and an open understory. We did use a tree-canopy percent-cover dataset in our modeling, but this was a generalized dataset and did not contain sufficient understory information.

Despite the limitations of our composite model and forecasting, this study provides the framework for understanding important habitat variables and how these variables might be affected in the future for forests similar to those on the HLCNF. Forest structure and prey are both important components of goshawk nest site locations. Although this paper did not cover prey densities we did analyze prey densities in this forest and for this system prey densities diffrences do not appear to be the reason for nest location (Wright et al. in preperation). The tools used in the paper are very usefull at identifying large scale forest structure changes. We understand that smaller scale forest structure changes are very important to understaand nest site locations. We would suggest in future work to start using using remnote sensing tools such as LIDAR to get small forest structure change data. Given the importance of goshawks as top-tier avian predators, this species remains an important focal point for the U.S. Forest Service in attempts to design management plans that conserve and maintain healthy forest ecosystems. Additionally, the distribution and unique nest site choices of goshawks provides the opportunity to use this species as an indicator of forest changes throughout the range. In northern forest areas, using goshawks to track forest changes gives us a powerful management tool to help better understand changing forest ecology, and address impacts.

## Acknowledgments

We would like to thank the Lewis and Clark National Forest. Without the on the ground support of Victor Murphy this project would not have been completed. This material is based upon work supported by NASA through contract NNL11AA00B and cooperative agreement NNX14AB60A.

Any opinions, findings, and conclusions or recommendations expressed in this material are those of the author(s) and do not necessarily reflect the views of the National Aeronautics and Space Administration (NASA)

